# A Fast 3 × *N* Matrix Multiply Routine for Calculation of Protein RMSD

**DOI:** 10.1101/008631

**Authors:** Imran S. Haque, Kyle A. Beauchamp, Vijay S. Pande

**Affiliations:** Stanford University, Departments of Computer Science, Counsyl, South San Francisco, CA; Stanford University, Departments of Biophysics Program Computational Biology Program, Memorial Sloan-Kettering Cancer Center, New York, NY; Stanford University, Departments of Chemistry

## Abstract

The bottleneck for the rapid calculation of the root-mean-square deviation in atomic coordinates (RMSD) between pairs of protein structures for large numbers of conformations is the evaluation of a 3 × *N* × *N* × 3 matrix product over conformation pairs. Here we describe two matrix multiply routines specialized for the 3 × *N* case that are able to significantly outperform (by up to 3X) off-the-shelf high-performance linear algebra libraries for this computation, reaching machine limits on performance. The routines are implemented in C and Python libraries, and are available at https://github.com/simtk/IRMSD.

## Introduction

The three-dimensional biomolecule structures provided by X-ray crystallography and NMR spectroscopy have pushed biology to the resolution of individual atoms. Understanding at this level of detail has been nothing short of revolutionary, with applications ranging from the design of small molecule drugs [8] to the detailed mechanism of protein synthesis within the cell [1, 13]. When analyzing biological structures, the root mean square deviation (RMSD) is a fundamental computational tool–the RMSD provides a valuable metric of the similarity of two conformations of a given protein. Such a metric enables algorithms that classify [10] protein structures and predict [12] structures from previously existing models.

RMSD plays an even larger role in analyzing molecular dynamics simulations of biomolecules. For example, Markov state model approaches [15, 4, 14] model the biomolecule dynamics by a Markov process on a state space that is typically determined by clustering. In such approaches, one often must cluster a simulation dataset with more than 10^6^ conformations into a model containing 10^4^ conformational states–clustering such a dataset is typically limited by the large number of RMSD calculations required.

Efficient, numerically stable algorithms exist to calculate both the structural RMSD as well as the optimal rotation matrix corresponding to this RMSD. Given some pre-calculated data, the RMSD can be calculated in 121 FLOPs (floating-point operations) on average [16], and the rotation matrix (as a unit quaternion) in 165 FLOPs in the worst case [11]. However, the pre-calculation steps for each algorithm are far more expensive. Both methods require the structures to be zero centered and require matrix magnitudes (self-inner-products) for both structures (where the coordinates are considered as 3×*N* matrices), as well as the inner product between the two. In large-scale clustering methods, the centering and magnitudes may be pre-calculated for each structure; however, the pairwise computation of coordinate matrix products takes approximately 18*N* FLOPs per matrix multiplication. Thus, even for small structures, the matrix multiplication becomes the bottleneck in RMSD calculation.

In this article we present a fast matrix multiply routine, specialized for the 3 × *N* matrix multiplications used in protein RMSD calculation, that uses SSE2/3 vectorization and register blocking for Intel and AMD x86_64 processors to run (3X) faster than off-the-shelf linear algebra routines. We present two versions of the matrix multiply, corresponding to different memory layouts, and provide benchmark results on internal protein datasets. Our code is available at https://github.com/simtk/IRMSD.

## 2 RMSD Calculation with the Theobald Algorithm

An algorithm due to Theobald [16] computes the RMSD between a pair of N-atom structures A and B (represented as *N* × 3 matrices) as a simple function of the largest eigenvalue of a 4×4 symmetric “key” matrix K: *RMSD* = [(*G_A_* + *G_B_* – 2*λ_max_*)*/N*]^1^*^/^*^2^. *G_A_* and *G_B_* are the self-inner-products of structures A and B (represented as *N* × 3 matrices), and the key matrix is computable from the elements of the product between A and B:

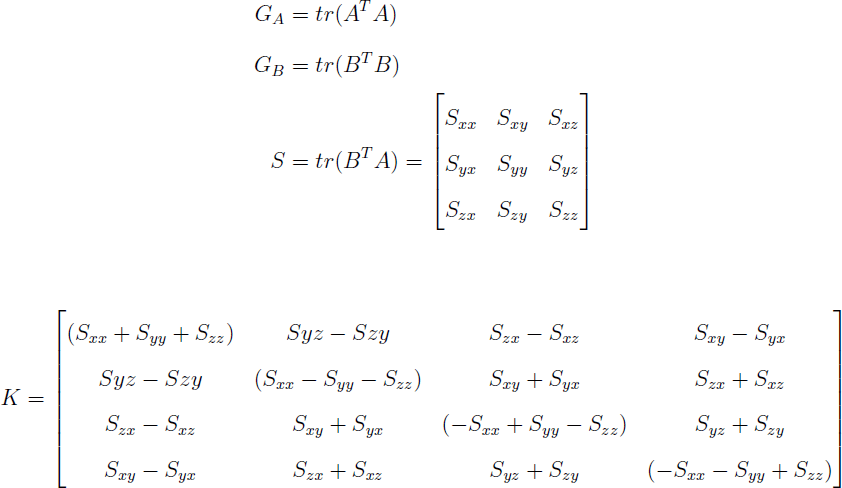

Given *G_A_*, *G_B_*, and *S*, Theobald’s algorithm, using Newton-Raphson iterations to refine the estimated eigenvalue, can compute the RMSD in 66 FLOPs (floating-point operations) to set up *K*, and an average of 55 FLOPs to iteratively estimate the eigenvalue. A refinement of the algorithm [11] also calculates the rotation matrix corresponding to the estimated RMSD in a worst-case 99 FLOPs after *K* has been computed.

## 3 Fast Matrix Multiplication Routines

Using the Theobald algorithm, the calculation of S dominates the cost of computing an RMSD. The *N* × 3 matrix inner product requires 9 × (*N* multiplies and (*N* – 1) adds); furthermore, it is memory-bandwidth intensive, requiring that both structures be loaded from memory at least once (three times, for a naive matrix multiply). Indeed, the matrix multiply is typically memory bandwidth bound.

In this section, we describe two versions of a register-blocked, vectorized matrix multiply to accelerate this computational bottleneck. Our target architecture is Intel/AMD x86_64 processors supporting the SSE vector intrinsics, which are currently the most common types of CPU found on standalone computers as well as large supercomputers. 64-bit support is helpful for these algorithms because it exposes an additional 8 architectural vector registers; our method can be used with only slight modification on 32-bit machines but will be less efficient because of register spills.

x86_64 SSE exposes a 4-wide (for single-precision) or 2-wide (for double-precision) vector architecture with 16 architectural vector registers. Modern CPUs (since the Core 2 generation from Intel and the K10 (Phenom) architecture from AMD) are able to execute a vector multiply or add in 1 clock cycle - potentially increasing throughput 4x from scalar floating point code. Our paper describes single-precision code. We have two different matrix multiplication routines, corresponding to the two common representations for protein structures in memory: axis-major and atom-major (Figure 1).

**Figure 1:**
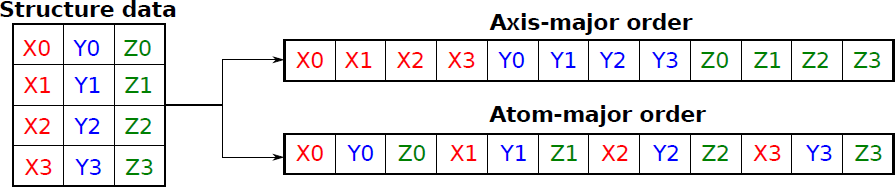
Illustration of linear layout options for a protein structure in memory. Xi/Yi/Zi are the X, Y, and Z coordinates of the i’th atom.

### 3.1 Axis-major Data

The simpler version of the matrix multiply requires that data be transposed from the representation used in the Theobald papers: specifically, instead of storing *N* × 3 matrices *A* and *B*, we store 3 × *N* matrices *A^T^* and *B^T^* . This “axis-major” layout has the x, y, and z axes along the slowly-varying index component of the matrix. These structures are stored such that each axis is padded with zeros such that the length is a multiple of 4 (to pack into vector registers properly), and each axis is stored in memory aligned to a 16-byte boundary (to use fast SSE aligned load instructions). In the description that follows, we will treat each axis of each structure as a 1 × *N* array *Ax*, *Ay*, etc.

The axis-major method performs *L/*4 iterations (where *L* is *N* rounded up to the nearest multiple of 4). Before the loop, 9 vector registers *xx*, *xy*, …, *zz* corresponding to the 9 elements of matrix *S* are initialized to [0,0,0,0]; these will hold partial summations of the axis-wise inner products. In each iteration, the method first loads 4 elements from each of *Bx*, *By*, and *Bz* into registers and then copied into temporary registers *tx*, *ty*, *tz*. It then loads 4 elements from *Ax* into a register. Since SSE has no non-destructive multiplication or addition operations, a destructive multiply is used to compute *tx = tx ∗ Ax*, and so on for each axis. This is then added to the accumulation register: *xx = xx* + *tx*. This set of operations then repeats for *Ay* and *Az*. In total, the loop as written uses all 16 vector registers in SSE: 9 accumulation registers, 3 to store copies of *Bx*, *y*, *z*, 3 temporaries, and 1 to store the loaded axis chunk from structure *A* (compiler optimization reduces the register count to 15). At the end of the loop, each vector accumulation register is then sum-reduced to a single element, corresponding to the required element of matrix *S*.

This method streams each structure through from memory only once (a factor of 3 improvement versus a naive multiply). On a 64-bit machine, moreover, it requires no memory accesses in the inner loop except for the structure loads; all temporaries are stored in registers, something made possible by the specialization to *N* × 3 matrices. Furthermore, with the exception of padding elements at the end and the final horizontal reduction (negligible in number for typical structure sizes), the vector lanes are fully utilized, dramatically improving arithmetic efficiency.

### 3.2 Atom-major Data

If storing data atom-major (i.e., as *N* × 3 rather than 3 × *N* in memory) is preferred for other reasons, it is still possible to gain many of the advantages of the previous vectorized method. Our atom-major method loads four atoms of coordinates from memory and shuffles the contents of the registers to recover an axis-major ordering in registers. This allows the same core arithmetic routines to be used as with the axis-major method, with an extra shuffle step at each load. Consequently, it eliminates the need for an expensive transpose operation in memory, at the cost of additional logic operations during the matrix multiplication. However, because the multi-core matrix multiply is typically memory-bound, these logic operations are often effectively “free” (hidden behind the latency of memory loads).

Figure 2 shows the structure of the shuffling operations used to transpose the atomic coordinates in registers. The temporaries needed for shuffling inflate the register count in the inner loop such that one register must be spilled to the stack each iteration. However, because this will almost certainly be served by the L1 cache, the performance impact is minimal. The four atoms (twelve coordinates) are first loaded into registers X, Y, and T2; Z is initialized to the value of X and T2 to the value of Y. The shuffle operation uses six instructions to unpack the data from X, Y, and T2 into registers X, Y, and Z, such that X contains the *x* coordinates for each of the 4 atoms, in order, and so on for Y and Z.

**Figure 2:**
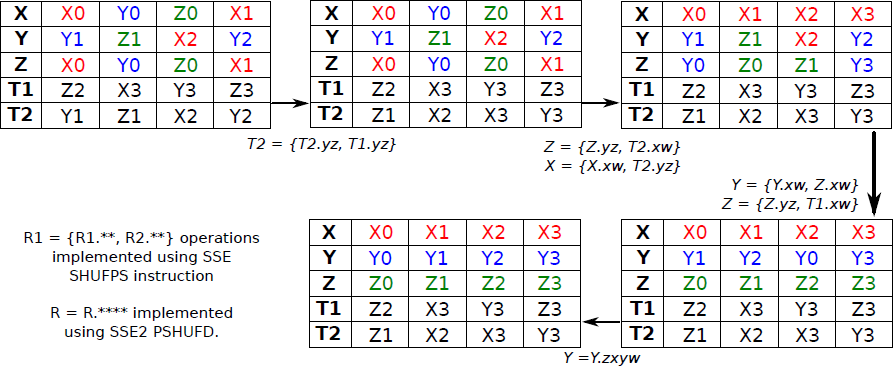
Dataflow for in-register transpose of atom-major data to axis-major format

## 4 Results

### 4.1 Benchmarking Methodology

To benchmark the performance of our matrix multiplication code, we compare its throughput to that of the matrix multiplication code included with the Theobald RMSD routines [16], as well as that of the SGEMM (single-precision matrix-matrix multiply) routine in GotoBLAS2 version 1.13 [7], a high-performance linear algebra library. We additionally compare the effective RMSD throughput of our matrix multiply and the GotoBLAS SGEMM as applied in k-centers clustering [6, 5] of protein conformations.

The matrix multiplication code was benchmarked on a machine with an Intel Core i7-920 CPU (2.66-2.93 GHz, 4 cores) with 12GB of DDR3-1066 memory (approx. memory bandwidth = 20 GB/s). All three matrix multiplication methods were compiled using gcc 4.3.3 with the following performance-relevant optimization flags: -O3 -funroll-loops -mtune=core2 -msse2 -msse3 -msse4. OpenMP, a multi-vendor language extension for parallelism in C/C++ and FORTRAN, was used to parallelize all matrix multiplication routines. The matrix multiplication benchmark first initializes 4 GB of random single-precision floating point numbers (2^30^ numbers). For each tested number of atoms *N,* it interprets this array as 2^30^*/N* structures of *N* atoms each. The first structure is treated as a reference, and every other structure is multiplied against this reference. For parallel runs, each thread handles a disjoint set of structures. This is run five times at each structure size and number of threads; the average of the elapsed time of the five runs is used to compute the effective arithmetic throughput of each method in GFLOP/s (billions of floating-point operations per second).

The clustering benchmark was performed using MSMBuilder [3] version 2.0. MSMBuilder contains a Python-Numpy implementation of the k-centers [6, 5] clustering algorithm using the Theobald RMSD metric. We modified MSMBuilder to use either the current multiplication code or GotoBLAS. The benchmark consisted of the PDBs 1NOT (176 atoms), 1YRF (582 atoms), 2KT2 (982 atoms), 2L30 (1679 atoms), 2KTD (2588 atoms), and 3PB4 (4947 atoms). This range of sizes covers all single-domain proteins with structures in the Protein Data Bank [2]. To generate nonequivalent protein conformations, we used the Gromacs [9] molecular dynamics package to simulate each protein for 5 ns. Clustering benchmarks were performed using single-threaded linear algebra. The K-Centers algorithm was used to cluster each dataset of 40,000 conformations into 100 states. The time required for clustering each dataset was measured using the Python `time.time()` function. Each benchmark was repeated 4 times; mean calculation times are reported. The time required for system preparation and file IO was subtracted; these values are independent of the underlying linear algebra library. Clustering benchmarks were performed on a 6-core AMD Phenom II X6 1090 T processor (3.2-3.4 GHZ) with 8GB of DDR3-1333 memory. The included benchmark reports single-threaded performance; as with the matrix multiplication benchmark, parallelism for both GotoBLAS and our code can be implemented using OpenMP to parallelize over structures.

### 4.2 Matrix Multiplication Throughput

The best way to evaluate the throughput of the methods is versus the architectural limits of our benchmark machine. The 20GB/s memory bandwidth is capable of sustaining 30 GFLOP/s in matrix multiplication if the reference structure is kept in the cache across comparisons or 15 GFLOP/s if both are streamed from memory. Each core of our Core i7-920 has a clock speed between 2.66 and 2.93 GHz (the CPU is capable of dynamically increasing its clock rate if thermal headroom is available), and can process 4 FLOP per cycle if instructions are issued one at a time (“single-issue throughput”), for an arithmetic peak of 10.64-11.72 GFLOP/s per core, or 42.56-46.88 GFLOP/s over all four cores. Thus, an optimal matrix multiplication code should achieve 10-12 GFLOP/s single-threaded (arithmetic-bound) and *∼* 30 GFLOP/s with four threads (memory-bound).

Figure 3 shows the performance of our code, GotoBLAS, and the original Theobald method with one and four threads. Four performance regimes are clearly visible for our code, corresponding to levels in the machine’s memory hierarchy (atom counts for which the reference structure can fit in the L1, L2, and L3 caches, and where it must be streamed from memory). With one thread, our axis-major code has the highest performance at all structure sizes; for small structures under 1,024 atoms, it is three times faster than the Theobald code and over twice as fast as GotoBLAS. For a small range of structure sizes (around 2,048 atoms), the axis-major method actually *exceeds* the machine’s single-issue peak. This is possible because the Core i7 CPU has separate pipelines for addition and multiplication; some of the operations in the inner loop can be issued simultaneously, thus allowing more than 4 FLOPs to complete per cycle. The extra shuffling logic in the atom-major method limits its performance in the single-threaded case; however, it still outperforms GotoBLAS by nearly a factor of two.

**Figure 3:**
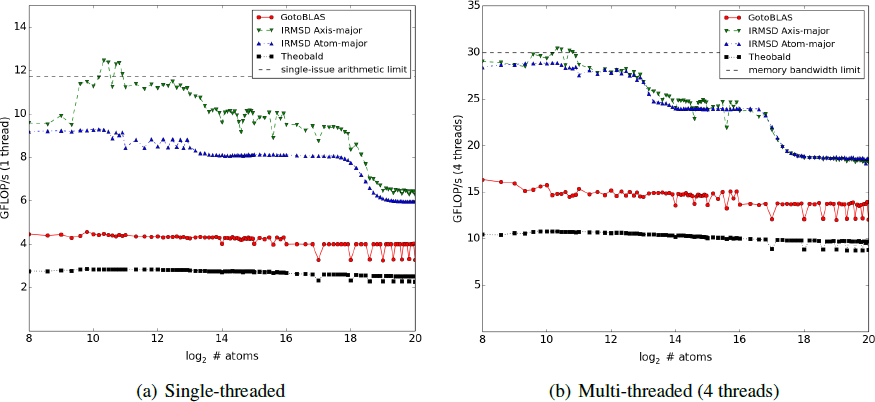
Arithmetic throughput of our matrix multiply routines versus GotoBLAS and matrix multiply from Theobald code in Ref [16], using 1 or 4 threads

The second half of Figure 3 illustrates the parallel performance of the various matrix multiplication methods. In this case, we expect all methods to be limited by the memory bandwidth of our test machine to around 30 GFLOP/s. Both the axis-and atom-major methods reach this limit for small structures. For larger structures that cannot fit in the L3 cache (right side of the graph), both methods still exceed 15 GFLOP/s, indicating that the threads are able to share some of the reference structure in cache, rather than reloading it individually. The performance of both of our methods is nearly identical, demonstrating that the extra shuffling logic of the atom-major method is irrelevant when throughput is memory bound. Both methods are approximately twice as fast as GotoBLAS and three times as fast as the original Theobald code.

### 4.3 Clustering Throughput

Table 1 shows the application-level improvement from using our matrix multiply method rather than an off-the-shelf linear algebra package. Protein clustering code using the axis-major method outperforms the same code using GotoBLAS by a factor of two to three.

**Table 1.** Clustering benchmark: speedup of axis-major method vs GotoBLAS

| PDB code | # atoms | Clustering time<br>with GotoBLAS (s) | Clustering time<br>with axis-major (s) | Speedup vs.<br>GotoBLAS |
| --- | --- | --- | --- | --- |
| 1NOT | 176 | 4.46 | 2.31 | 1.93 × |
| 1YRF | 582 | 15.24 | 5.87 | 2.60 × |
| 2KT2 | 982 | 25.67 | 8.43 | 3.05 × |
| 2L30 | 1679 | 41.77 | 12.64 | 3.30 × |
| 2KTD | 2588 | 61.29 | 21.31 | 2.88 × |
| 3PB4 | 4947 | 118.89 | 37.70 | 3.15 × |

## 5 Conclusions

Existing linear algebra libraries cannot efficiently handle the very narrow-aspect-ratio matrix multiplications required for efficient calculation of protein RMSD. We have described vectorized methods for matrix multiplication on the x86_64 architecture that provide a factor of two to three speedup relative to current high-speed linear algebra packages. Our routines are able to saturate machine limits for throughput, significantly accelerating protein RMSD calculations for large-scale clustering and protein search applications. The routines are implemented in C and Python libraries, and are available at https://github.com/simtk/IRMSD.

## Acknowledgements

The authors thank John Chodera, Peter Kasson, and Kai Kohlhoff for their implementations of the Theobald algorithm and Greg Bowman for protein structure data for benchmarking and testing. We thank NSF CNS-0619926 for computing resources and NIH R01-GM062868, NSF-DMS-0900700, and NSF-MCB-0954714 for funding.

